# The Human Kinesin Kif18A’s Neck Linker Permits Navigation of Microtubule Bound Obstacles within the Mitotic Spindle

**DOI:** 10.1101/364380

**Authors:** Heidi L. H. Malaby, Dominique V. Lessard, Christopher L. Berger, Jason Stumpff

**Affiliations:** Department of Molecular Physiology and Biophysics, University of Vermont, Burlington, VT 05405, United States

**Keywords:** kinesin, Kif18A, mitosis, HURP

## Abstract

Mitotic chromosome alignment is essential for the robust separation of genetic material into daughter cells. In mammalian cells, this process requires the function of Kif18A, a kinesin-8 motor protein. Kif18A confines chromosome movement to the mitotic spindle equator by accumulating at the plus-ends of kinetochore microtubule bundles (K-fibers), where it functions to suppress K-fiber dynamics. It is not understood how the motor accumulates at K-fiber plus-ends, a difficult feat requiring the motor to navigate protein dense microtubule tracks. Our data indicate that Kif18A’s relatively long (17 amino acid) neck linker is required for the motor’s accumulation at K-fiber plus-ends. Shorter neck linker (sNL) variants of Kif18A display a deficiency in K-fiber accumulation, especially on K-fibers near the center of the spindle. This pattern correlates with the more uniform concentration of the microtubule bundling protein HURP on central K-fibers compared to peripheral K-fibers. Depletion of HURP permits Kif18A sNL to accumulate on central K-fibers, while HURP overexpression reduces wild-type Kif18A’s ability to accumulate on this same K-fiber subset. Furthermore, single molecule assays indicate that Kif18A sNL motors are less proficient at navigating microtubules coated with the microtubule associated protein tau. Taken together, these results support a model in which Kif18A’s neck linker length permits efficient navigation of obstacles such as HURP to reach K-fiber ends during mitosis.

**Signficiance Statement:** Kinesin motor proteins play key roles in controlling chromosome alignment and segregation during cell division. The kinesin Kif18A confines chromosomes to the middle of the spindle by accumulating at the ends of microtubules attached to chromosomes. We show here that Kif18A’s ability to accumulate at the end of these microtubules requires navigation of microtubule-associated protein obstacles, and that this activity is imparted by a relatively long neck linker region. These findings demonstrate a molecular mechanism for navigation of densely populated microtubules inside a cell.

## Introduction

Kinesin motor proteins are responsible for building and maintaining the mitotic spindle (1, 2), transporting, aligning, and orienting chromosomes at the metaphase plate (3-5), and scaffolding the spindle midzone for cytokinesis (6). These microtubule-dependent functions must be performed in the context of dense molecular environments due to the sheer number of microtubule associated proteins (MAPs) that are found within mitotic spindles (7).

The kinesin-8 motor protein, Kif18A, functions in this environment to confine chromosome movements around the metaphase plate and is required for proper chromosome alignment (4, 8). Kif18A accumulates at the ends of K-fibers, which are bundles of ~ 15-20 microtubules collectively bound to kinetochore protein complexes assembled at centromeres (9). Kif18A accumulation on microtubule ends suppresses microtubule dynamics (4, 10), dampening chromosome oscillations in metaphase (11).

Kif18A is a dimeric, processive kinesin, containing two conserved globular motor domains that interact with microtubules through surface polar and positively charged residues (12). Like all ATP dependent kinesins, motor domain affinity for the microtubule surface correlates with nucleotide state. The neck linker, a short 14-17 amino acid region between the end of the motor domain and the coiled-coil stalk, begins to dock to the motor domain upon a conformational shift induced by ATP binding, and finishes docking upon hydrolysis of ATP to ADP and phosphate release (13), creating a lever action to pull forward the ADP-bound trailing motor domain (14). While this mechanical process is highly conserved among members of the kinesin family, small residue differences can alter the balance between processivity and microtubule affinity, determining the motor off rate (14). For example, increasing the neck linker length of Kinesin-1 (*Drosophila* conventional, normally 14 residues) reduces its run length, while shortening the Kinesin-2 neck linker (Kif3A, normally 17 residues) enhances its run length (15). In addition, the longer neck linker length of Kinesin-2 provides the structural flexibility required to navigate around microtubule-bound obstacles (16-18). These studies have all been performed in purified *in vitro* systems. The importance of neck linker flexibility for kinesin motility in cells is not understood.

While a C-terminal microtubule binding domain is necessary for Kif18A accumulation on K-fibers in cells (19-21), it is not sufficient. A Kinesin-1 chimera containing Kif18A’s C-terminus does not accumulate on K-fiber ends, indicating there may be additional structural determinants of Kif18A allowing for K-fiber end accumulation (22). Here, we determined if Kif18A’s neck linker length, which is 17 residues, is important for its ability to accumulate at K-fiber ends. We created a panel of Kif18A short neck linker (sNL) constructs and show that all of these are deficient in accumulation on K-fibers at the center of the spindle. These sNL Kif18A constructs are deficient for chromosome alignment and promoting progression through mitosis. In a purified system, we show that shortening the Kif18A neck linker creates a faster, less processive motor that is not as efficient at navigating obstacles on the microtubule. Importantly, we identify the microtubule-bundling protein HURP as one obstacle of Kif18A in mitotic cells, and show that reduced expression of HURP allows Kif18A sNL to accumulate on central K-fibers, while increased HURP expression inhibits accumulation of the wild-type motor. Taken together, this study supports a model in which Kif18A’s long neck linker tunes the motor for navigation of the densely populated K-fiber surface.

## Results

### Shorter neck linker variants of Kif18A do not accumulate at the ends of K-fibers in the center of the spindle

To investigate the importance of Kif18A neck linker length on its ability to accumulate on K-fiber plus-ends in mitotic cells, we created a panel of shorter neck linker (sNL) Kif18A variants (Figure 1). To determine where to make these deletions, we used a combination of coiled-coil prediction algorithms (COILS and LOGICOIL) and sequence alignments with Kif5B and Kif3A. From these we concluded that Kif18A’s probable neck linker is 17 amino acids long from residues 353-369. Because coiled-coil prediction algorithms are inaccurate at pinpointing the beginning of coiled-coil domains and altering sequences in kinesin neck linkers can affect the coiled-coil (23), we created a series of four short neck linker variants, taking care to keep the highly conserved N362 in frame (sNL0, sNL1, sNL2, sNL3 Figure 1B). To test the localization of these constructs on K-fibers, we treated HeLa cells with Kif18A siRNA, then transfected each GFP-Kif18A sNL variant, which also contained silent mutations in the siRNA-targeted region. Cells were arrested in metaphase by treatment with MG132, fixed, and stained with a GFP antibody. GFP-Kif18A sNL0 showed no K-fiber end accumulation and was unable to align chromosomes, suggesting that it may be inactive (Figure S1A-B). We further tested activity by treating cells with control or Kif18A siRNAs, then transfecting GFP-Kif18A sNL0. Before fixation, cells were treated with taxol for 10 minutes, which stabilizes K-fibers and promotes end accumulation (22). When endogenous Kif18A is present, the majority of cells (~70%) show some GFP-Kif18A sNL0 accumulation on K-fibers; however, this accumulation is diminished (< 20%) when the endogenous motor is knocked down (Figure S1C). This finding suggests that Kif18A sNL0 is only active when heterodimerized with endogenous Kif18A. Thus, we did not pursue this construct further.

**Figure 1.**
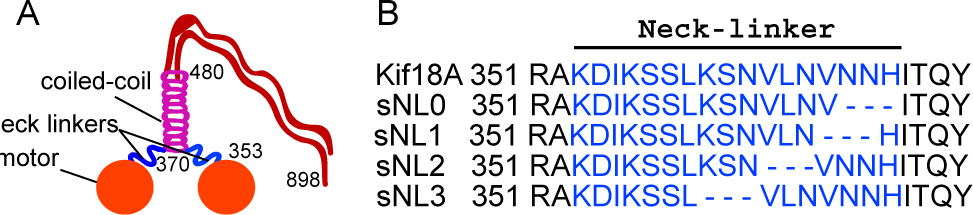
Kif18A putative structure and predicted neck linker region. (**A**) Cartoon depiction of known Kif18A structural regions with residue numbers. (**B**) Predicted neck linker residues (blue) of Kif18A and short neck linker (sNL) variants used in this study.

The localizations of GFP-Kif18A sNL1, sNL2, and sNL3 in mitotic cells were similar and differed from the uniform distribution exhibited by sNL0 (Figure 2). Strikingly, none of these Kif18A sNL variants accumulated on K-fibers at the center of the spindle (Figure 2A, white arrows) but did accumulate on K-fibers near the spindle periphery (Figure 2A, blue arrows and Figure S2). We quantified the localization of each sNL mutant via line scan analysis, averaging multiple profiles from many cells by aligning to the peak intensity of anti-centromere antibody (ACA) immunofluorescence, which labels kinetochores. The lack of accumulation for any GFP-Kif18A sNL variant in the central spindle was a highly reproducible effect (Figure 2B). These results indicate that 1) Kif18A sNL1-3, but not sNL0, are active motors and 2) a longer neck linker is necessary for central spindle K-fiber accumulation in mitotic cells.

**Figure 2.**
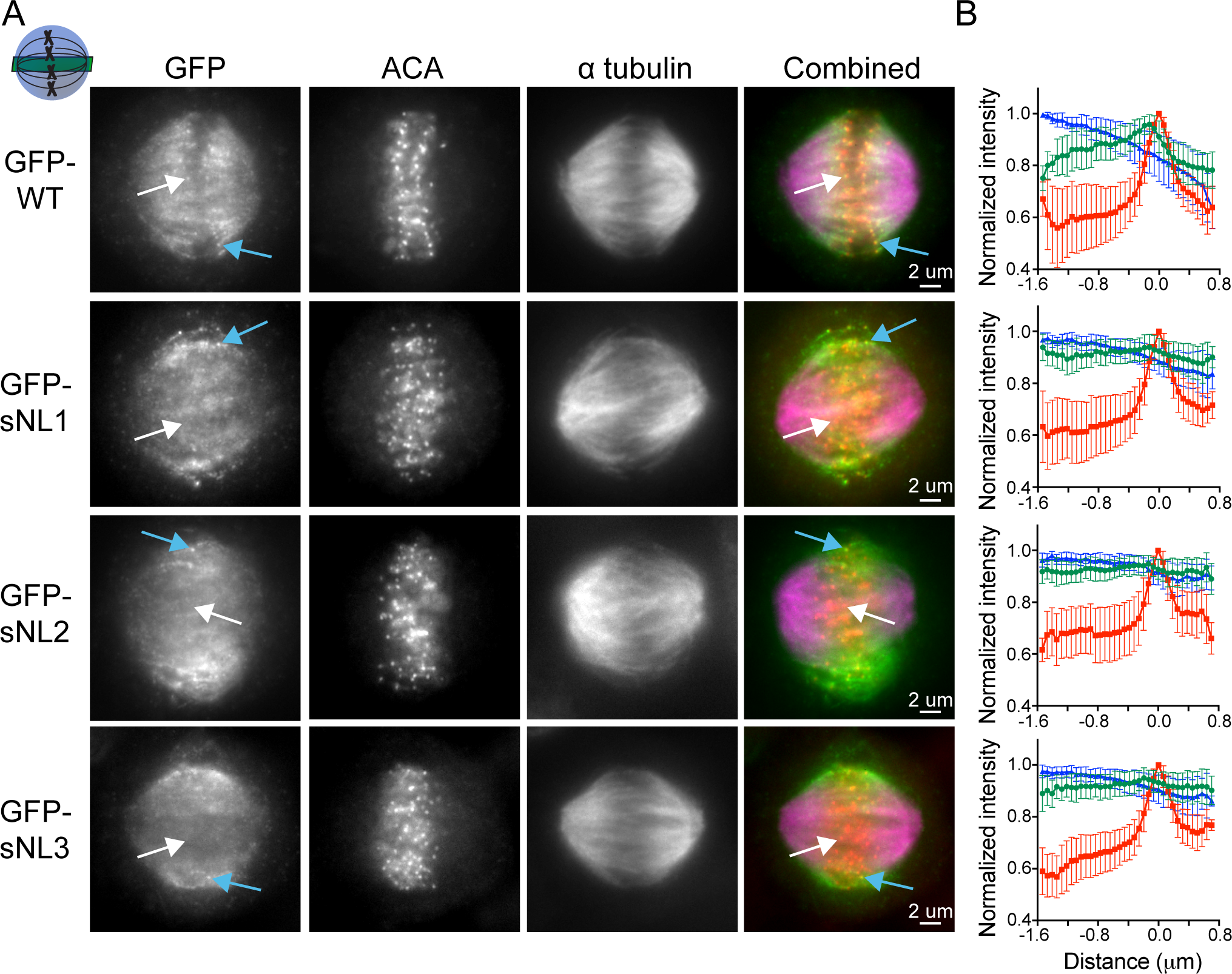
Kif18A sNL variants do not accumulate on K-fibers in the center of the spindle. (**A**) Representative metaphase arrested cells expressing GFP-WT Kif18A or GFP-sNL1-3 variants. Shown is one z-slice through the middle of the spindle. White arrows point to a central K-fiber in each cell, blue arrows point to a peripheral K-fiber. (**B**) Quantification of Kif18A accumulation on central K-fiber ends by line scan analysis. Individual lines were normalized and aligned to peak ACA intensity. Symbols show normalized intensity average, error bars are standard deviation. Red, ACA. Blue, a-tubulin. Green, Kif18A variant. p < 0.0001 by F test comparing slopes of linear regressions for WT vs. any sNL variant. Data obtained from two independent experiments with the following cell and line numbers: GFP-WT (13 cells, 40 lines), GFP-sNL1 (17 cells, 49 lines), GFP-sNL2 (15 cells, 51 lines), GFP-sNL3 (20 cells, 62 lines).

### Kif18A sNL variants are deficient in chromosome alignment and cause a mitotic delay

We next determined if this Kif18A sNL central spindle accumulation deficiency had any effect on the motor’s ability to align chromosomes. In metaphase cells lacking Kif18A, chromosomes and kinetochores are broadly distributed within the spindle (4, 22). We knocked down endogenous Kif18A in HeLa cells, then transfected each GFP-sNL Kif18A variant to measure its ability to align mitotic chromosomes. For comparison, we transfected Kif18A knockdown cells with GFP-WT Kif18A as a positive control and GFP alone as a negative control. Cells were arrested in metaphase, fixed, and stained for kinetochores, GFP, and γ-tubulin to label spindle poles (Figure 3A). To quantify alignment, we measured the distribution of kinetochore (ACA) immunofluorescence along the spindle axis and determined the full width at half maximum (FWHM) of this distribution (11, 22). As expected, GFP alone did not rescue chromosome alignment, while GFP-WT Kif18A fully recovered alignment (Figure 3B). All three GFP-sNL Kif18A variants produced intermediate chromosome alignment phenotypes, where often the chromosomes off the metaphase plate were in the central region of the spindle where the motor did not accumulate (white arrows). Kif18A expression is also known to affect spindle length, so we also quantified the pole-to-pole spindle length (SL) distance (Figure 3B). Again, the Kif18A sNL variants as a group displayed intermediate spindle lengths.

**Figure 3.**
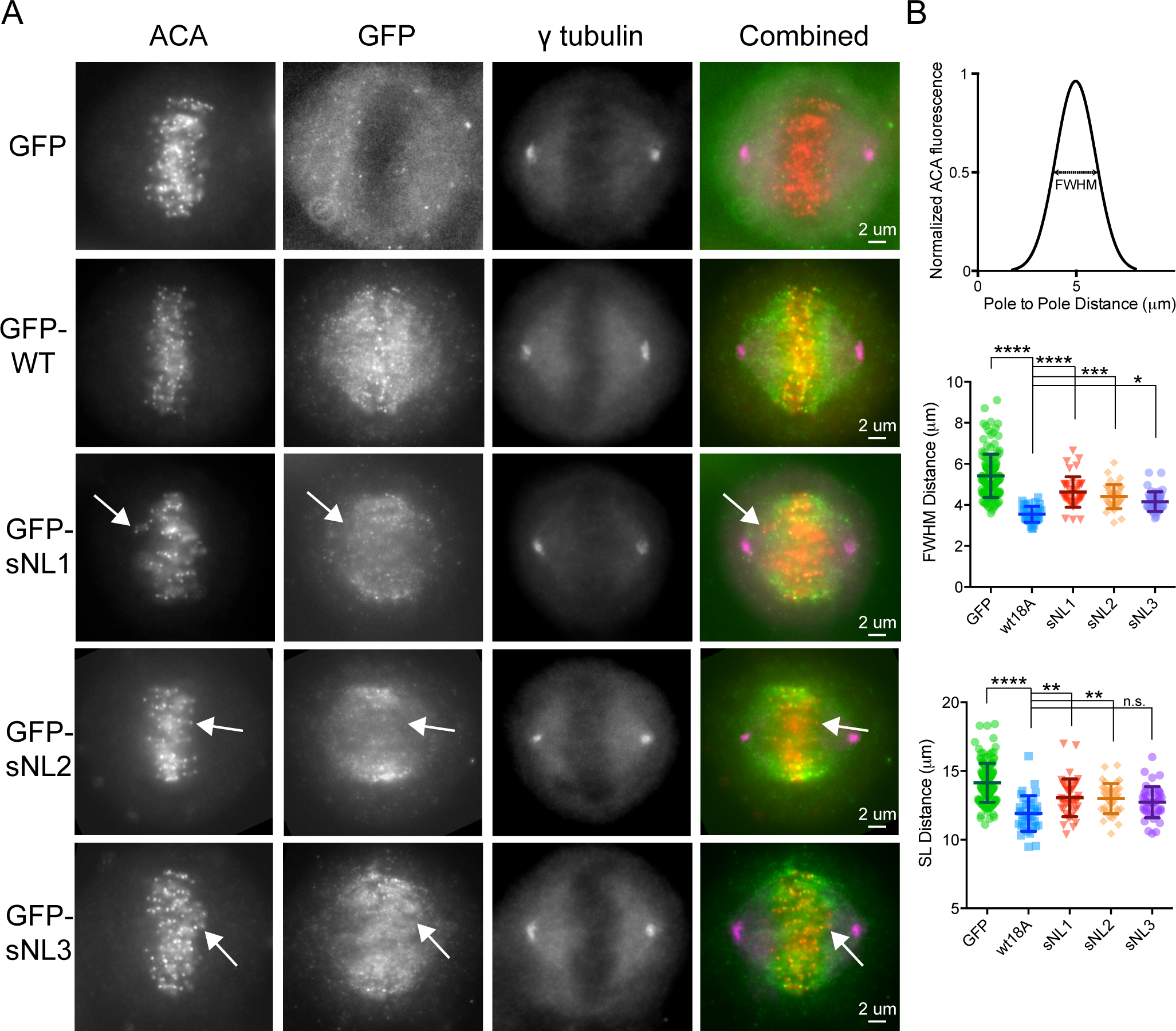
Kif18A sNL variants are deficient in chromosome alignment. (**A**) Representative metaphase arrested cell images where all cells were first treated with Kif18A siRNAs, then transfected with GFP alone, GFP-WT Kif18A or GFP-sNL1-3 variants. White arrows mark a chromosome pair off the metaphase plate in the central spindle where GFP-sNL variant is not accumulating. Images are one z-plane through the middle of the cell where both poles are in focus. (**B**) (*top*) Graphical explanation of ACA fluorescent signal intensity FWHM measurement. (*middle*) FWHM distance and (*bottom*) spindle length (SL) distance quantifications for all conditions. ****, adjusted p < 0.0001, ***, adjusted p = 0.0001, *, adjusted p = 0.0119, with 95% confidence interval by one-way ANOVA with Tukey’s multiple comparisons test. Data obtained from three independent experiments with the following cell numbers: GFP only (141), GFP-WT Kif18A (35), GFP-sNL1 (41), GFP-sNL2 (38), GFP-sNL3 (43).

Loss of Kif18A function in HeLa cells also leads to an extended mitotic delay that is dependent on the spindle assembly checkpoint (8, 24). Control HeLa cells progress through mitosis in ~ 40 minutes, whereas cells deficient in Kif18A can be arrested for hours (25). Since Kif18A sNL1-3 variants all had similar localization and alignment characteristics, we focused the rest of our studies on Kif18A sNL1. Endogenous Kif18A was knocked down in HeLa cells, and cells were transfected with GFP-WT Kif18A, GFP-sNL1, or GFP alone (Figure 4A). Cells were analyzed by time-lapse microscopy and the time from nuclear envelope breakdown (NEB) to anaphase onset (AO) was determined for each GFP expressing cell that divided completely. GFP-sNL1 Kif18A displayed a ~ 20 minute mitotic delay over GFP-WT Kif18A (Figure 4B). Taken together, these findings show that Kif18A’s long neck linker is required for complete chromosome alignment and timely progression through mitosis.

**Figure 4.**
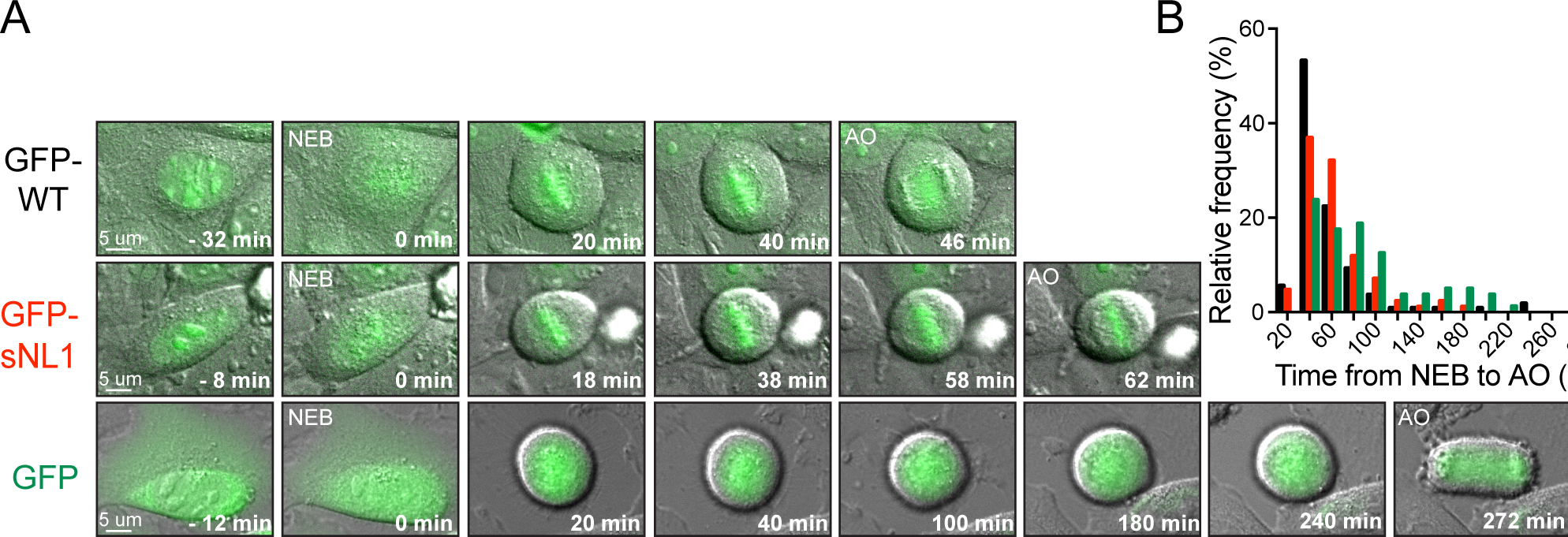
Kif18A sNL variant fails to promote proper mitotic progression. (**A**) Time course DIC and GFP overlay images from live cells treated with Kif18A siRNAs and then transfected with GFP-WT Kif18A, GFP-sNL1, or GFP alone. NEB, nuclear envelope breakdown. AO, anaphase onset. Times are normalized to NEB. (**B**) Histogram distributions of the time observed from NEB to AO for all cells with GFP expression that finished division by the end of imaging. For chi square analysis, all conditions were subdivided into 20 minute bins, then grouped into three timeframes: < 40 minutes, 60-80 minutes, > 100 minutes. GFP-WT Kif18A (expected) to GFP-sNL1 (observed): p = 0.0247; GFP-WT Kif18A (expected) to GFP alone: p < 0.0001; GFP alone (expected) to GFP-sNL1 (observed): p = 0.0006. Data obtained from three independent experiments with the following cell numbers: GFP-WT Kif18A (107), GFP-sNL1 (84). GFP alone (80).

### Kif18A sNL1 is faster than WT Kif18A but has a shorter run length

Neck linker length is an important determinant of run length and velocity for kinesin-1 and kinesin-2. For example, shortening the 17 residue Kinesin-2 neck linker increases both run length and velocity (15). To determine what effect shortening the Kif18A neck linker has on its motile properties, we compared purified Kif18A WT and sNL1 in single molecule assays. Because we wanted to observe only the neck linker’s influence on velocity and run length, we used truncated Kif18A constructs (Δ480) that lack the C-terminal microtubule-binding domain. The movement of individual GFP-tagged motors were visualized on rhodamine-labeled microtubules via TIRF microscopy (Figure 5A). The velocity and run length of GFP-Kif18A Δ480 was comparable to those measured for other truncated Kif18A constructs (19, 20). In contrast, GFP Kif18A Δ480 sNL1 was almost twice as fast as the WT motor, but moved about one third the distance (Figure 5B and 5C, Table S1). However, this decreased run length alone would not explain why Kif18A sNL variants are able to accumulate at the ends of peripheral but not central K-fibers, especially since peripheral K-fibers, assuming relative continuity, are predicted to be slightly longer due to the geometry of the spindle.

**Figure 5.**
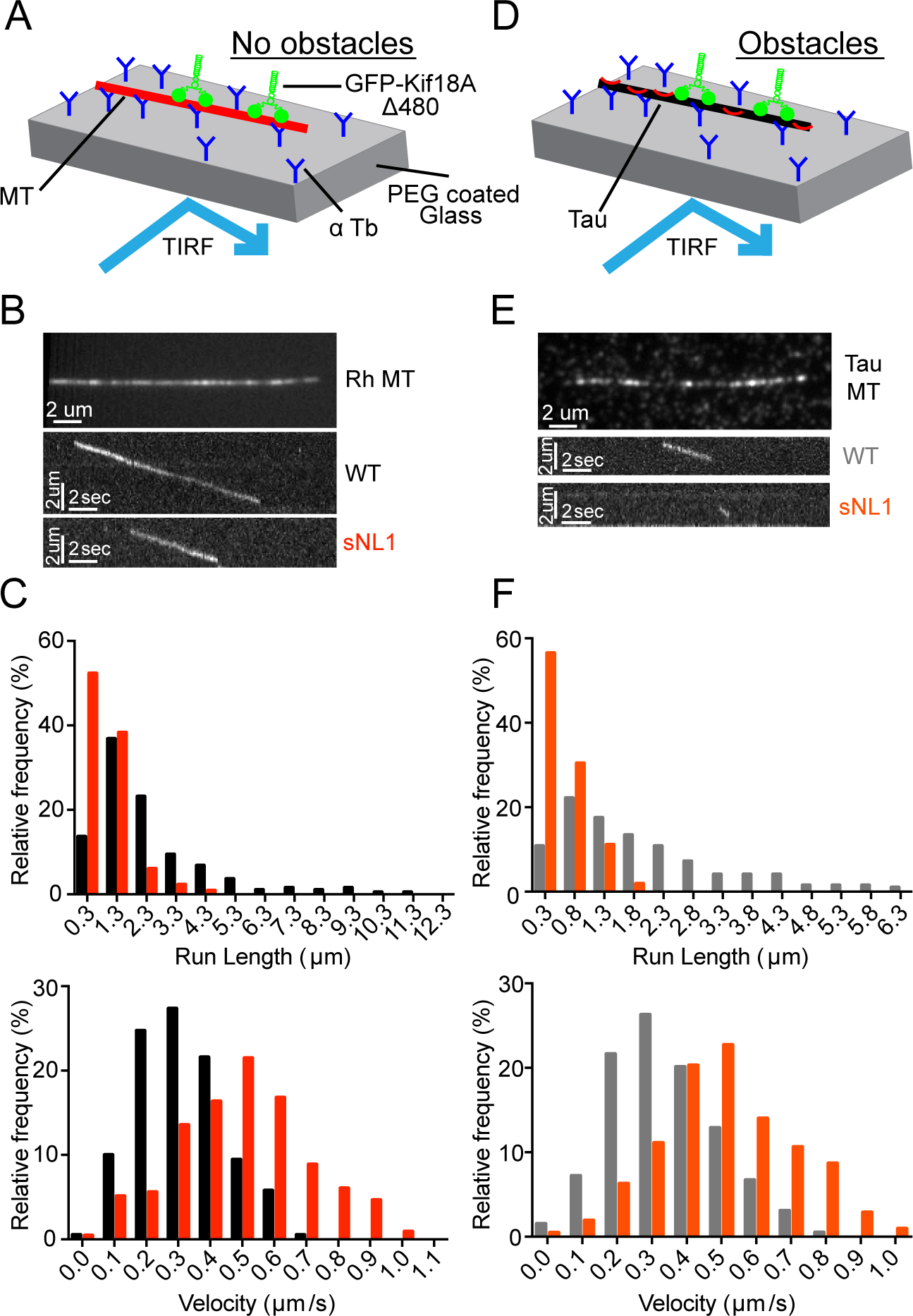
Kif18A sNL variant is less processive but faster than WT Kif18A and more hindered by obstacles. (**A**) Cartoon depiction of single molecule TIRF assay set up without obstacles on microtubules (MT). (**B**) (*top*) Picture of single Rhodamine (Rh) labeled microtubule in single molecule TIRF. (*bottom*) Representative kymographs of GFP-Kif18A Δ480 WT (WT) and GFP-Kif18A Δ480 sNL1 (sNL1) processive events. (**C**) Histogram distributions of run lengths (*top*) and velocities (*bottom*) of processive events observed. p < 0.0001 for both run length and velocity comparing WT and sNL1 by unpaired t-test. (**D**) Cartoon depiction of single molecule TIRF assay set up with Alexa 647 3RS tau as a diffusive obstacle on microtubules. (**E**) (*top*) Picture of single Alexa 647 3RS tau coated microtubule. (*bottom*) Representative kymographs of GFP-Kif18A Δ480 WT (WT) and GFP-Kif18A Δ480 sNL1 (sNL1) processive events on tau-coated microtubules. (**F**) Histogram distributions of run lengths (*top*) and velocities (*bottom*) of processive events observed on tau-coated microtubules. Statistical significance comparing no tau vs. tau by unpaired t-test: Run length, WT p = 0.0096; sNL1 p < 0.0001; Velocity, WT p = 0.0407; sNL1 p = 0.5926. Data obtained from two independent protein preps and 16 individual TIRF assays with the following total processive events: No tau: WT (190), sNL1 (214); With tau: WT (194), sNL1 (207).

### The motility of Kif18A sNL1 is more hindered in the presence of obstacles than WT Kif18A

The structural flexibility imparted by the long neck linker of kinesin-2 permits navigation around obstacles on microtubules (16, 18). To determine how Kif18A’s motility is affected by the presence of microtubule-associated proteins, we compared the movement of single GFP-Kif18A Δ480 WT molecules in the presence and absence of rigor kinesin and tau. Rigor kinesin was used at a concentration of 1:12.5 rigor kinesin to tubulin dimer, which limits the movement of kinesin-1 motors (18). The presence of rigor kinesin did not alter the total run length or average velocity of GFP-Kif18A Δ480 WT but did increase the frequency of motor pausing (Figure S3A-D). The microtubule associated protein tau binds both statically and diffusely to the microtubule surface (26, 27) and its presence reduces the run length of kinesin-1 (28-30). We found tau reduced the run length of GFP-Kif18A Δ480 WT by ~10%. However, tau’s effect on run length was significantly larger (over 50%) for GFP-Kif18A Δ480 sNL1 (Figure 5D-F and Table S1). Tau did not alter the velocity of either motor. Taken together, these findings indicate WT Kif18A is capable of obstacle navigation on microtubules, and shortening Kif18A’s neck linker significantly compromises this activity.

### HURP is an obstacle for Kif18A on the mitotic spindle

Our single molecule studies suggest that the localization and functional defects observed for Kif18A sNL1 in mitotic cells could be caused by an inability of the motor to navigate around microtubule associated proteins to reach the ends of K-fibers. The microtubule-stabilizing protein HURP regulates Kif18A’s localization and function in mitotic cells (31). HURP localizes as a gradient on K-fibers with a concentration near the plus-ends due to Ran regulation (32, 33). In HURP-depleted cells, Kif18A displays a tighter accumulation at K-fiber plus-ends, while overexpression of HURP’s microtubule-binding domain disrupts Kif18A’s localization and function. Interestingly, these latter effects can be rescued by overexpression of Kif18A (31). These results are consistent with HURP functioning as an obstacle or filter for motor movement to the K-fiber plus-end.

To test this idea, we treated HeLa cells with siRNAs against endogenous Kif18A or endogenous Kif18A and HURP, then transfected GFP-WT Kif18A or GFP-sNL1. Cells were arrested in metaphase, fixed, and stained for GFP, endogenous HURP, and kinetochores (ACA) (Figure 6A). To quantify Kif18A accumulation changes at K-fiber ends in the center of the spindle, we generated normalized, aligned line scans of GFP fluorescence as in Figure 1 and determined the area under the curve for K-fiber regions close to (tip) and farther from (lattice) the kinetochore. We computed the ratio of tip to lattice Kif18A fluorescence as a metric for its plus-end accumulation (Figure 6C). After HURP knockdown, GFP-WT Kif18A showed a slight decrease in central K-fiber accumulation, which we attribute to less microtubule tracks present per K-fiber bundle upon HURP knockdown (Figure S4) (34). Despite the decrease in K-fiber attachment and stability caused by HURP-depletion, GFP-sNL1 Kif18A displayed a significant increase in central spindle accumulation (Figure 6C, middle). Moreover, when we knocked down endogenous Kif18A and then co-expressed GFP-HURP with either mCherry-tagged Kif18A WT or sNL1 (Figure 6B), which led to stronger K-fiber immunofluorescence (Figure S4), WT Kif18A accumulation was reduced in the central spindle (Figure 6C, right).

**Figure 6.**
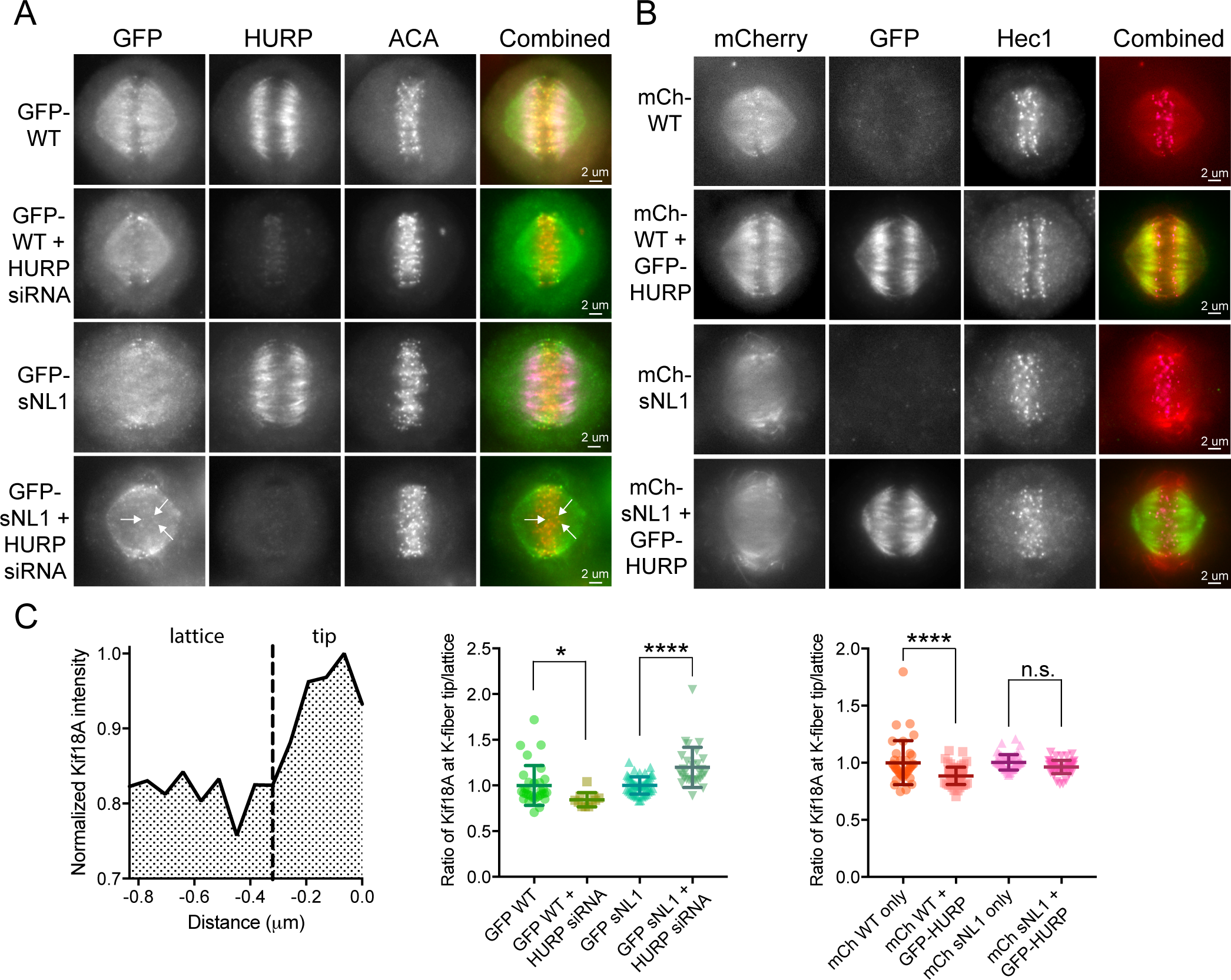
HURP is an obstacle for Kif18A on K-fibers in metaphase cells. (**A**) Metaphase arrested cells treated with Kif18A siRNA (and HURP siRNAs in marked conditions) and transfected with GFP-WT Kif18A or GFP-sNL1. (**B**) Metaphase arrested cell treated with Kif18A siRNA and transfected with mCherry-WT Kif18A, mCherry-sNL1, or GFP-HURP. Shown in (A) and (B) is one z-slice through the middle of the spindle. (**C**) Quantification of Kif18A accumulation at central K-fibers ends under the above conditions. (*left*) Line scans were performed, normalized, and aligned as in Figure 2, then each line was divided at the dotted line (−0.32 μm) and the area under the curve calculated for each segment. The ratio of tip (0 to ‐0.32 μ from kinetochore) to lattice (−0.32 μm to ‐0.768 μm) Kif18A fluorescent intensity is reported in the following graphs. (*middle*) Quantification of GFP-WT and GFP-sNL1 with HURP knockdown. *, adjusted p = 0.0429, ****, adjusted p < 0.0001 with 95% confidence interval by one-way ANOVA with Tukey’s multiple comparisons test. Data obtained from two independent experiments with the following cell and line numbers: GFP-WT (12 cells, 31 lines), GFP-WT + HURP siRNA (7 cells, 11 lines), GFP-sNL1 (14 cells, 50 lines). GFP-sNL1 + HURP siRNA (12 cells, 29 lines). (*right*) Quantification of mCherry-WT and mCherry-sNL1 with GFP-HURP overexpression. ****, adjusted p < 0.001, n.s., not significant (adjusted p = 0.2061) with 95% confidence interval by one-way ANOVA with Tukey’s multiple comparisons test. Data obtained from two independent experiments with the following cell numbers: mCh-WT (13 cells, 37 lines), mCh-WT + HURP-GFP (16 cells, 63 lines), mCh-sNL1 (12 cells, 48 lines). mCh-sNL1 + GFP-HURP (16 cells, 51 lines).

We also questioned how Kif18A sNL variants could accumulate on peripheral but not centrally located K-fibers within the spindle, as both sets of K-fibers appear to contain HURP. However, closer examination of HURP and Kif18A localization indicates that both WT and Kif18A sNL localize to zones of low HURP on peripheral K-fibers, suggesting that Kif18A sNL1 is able to accumulate on peripheral K-fibers because HURP is not uniformly present on these tracks (Figure S5). Taken together, these results indicate that the presences of HURP on K-fibers reduces Kif18A’s ability to accumulate at plus-ends and that this effect is exacerbated by shortening the Kif18A neck linker.

## Discussion

Our work supports a model in which Kif18A must exhibit agility during its translocation along K-fibers to navigate microtubule-associated proteins such as HURP and accumulate at plus-ends, where it functions to control chromosome movements (Figure 7). Kif18A’s relatively long 17 amino acid neck linker is an important molecular component underlying the motor’s navigation activity. Based on studies of the kinesin-1 and kinesin-2 neck linkers, we predict that Kif18A’s neck linker region provides structural flexibility, allowing the motor to step from one protofilament to the next as it moves towards microtubule plus-ends (18). In support of this idea, the budding yeast ortholog of Kif18A, Kip3p, which shares a highly conserved neck linker region, is capable of switching protofilaments during movement (35, 36). However, a physiological role for this type of side-stepping has not been determined for any kinesin. We propose that this lateral movement is necessary for Kif18A’s mitotic functions based on our observations that shorter neck linker variants of Kif18A 1) display hindered motility in the presence of microtubule-bound obstacles *in vitro* similar to kinesin-1 and kinesin-2 motors with short neck linkers (15-18), 2) are not able to accumulate properly on K-fibers in mitotic cells unless the microtubule-associated protein HURP is depleted, and 3) fail to align chromosomes or promote proper progression through mitosis in the absence of endogenous Kif18A.

**Figure 7.**
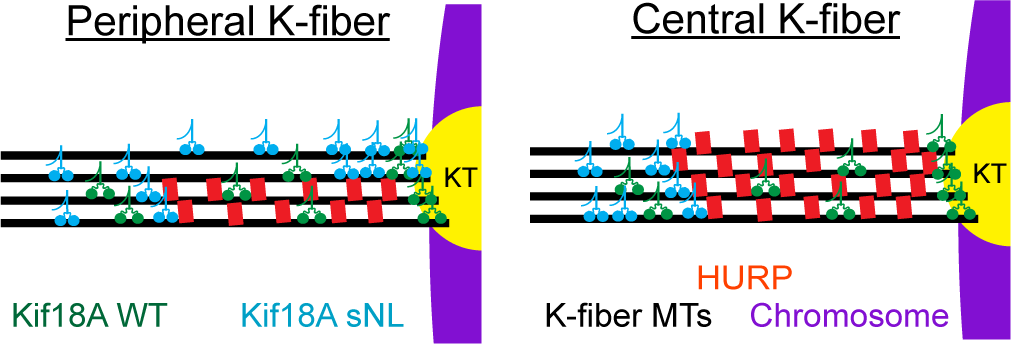
Model for the role of Kif18A’s neck linker in K-fiber end accumulation. Kif18A WT is ideally suited to navigate obstacles, like HURP, on K-fibers. Conversely, shortening Kif18A’s neck linker reduces this navigation ability and limits accumulation to the outside of peripheral K-fibers. KT, Kinetochore protein complex.

Our model that HURP acts as an obstacle for Kif18A is also consistent with previously published data showing that overexpression of HURP or the microtubule-binding domain of HURP inhibits Kif18A accumulation at K-fiber plus-ends (31). Increases or decreases in HURP expression directly affect the stability of microtubules in K-fibers, a finding that is true for other microtubule bundling proteins as well (37). HURP knockdown reduces K-fiber stability and WT Kif18A accumulation at K-fiber plus-ends. In contrast, Kif18A sNL1 accumulation at K-fiber plus-ends increases following HURP knockdown. This result is consistent with obstacles being removed and the potential of Kif18A sNL1 to reach the ends of less stable microtubules faster than WT Kif18A based on its increased velocity. On the other hand, HURP overexpression in cells stabilizes K-fibers but also reduces WT Kif18A accumulation at plus-ends. These results suggest that a balance of Kif18A and HURP expression must be maintained to ensure that chromosome movements are properly controlled and spindle stability is maintained.

Our single molecule studies indicate that Kif18A sNL1 is twice as fast as WT Kif18A, but only travels a third of the distance. In contrast, shortening Kif3A’s neck linker from 17 to 14 residues resulted in a faster motor that traveled longer distances (16), and lengthening Kinesin-1’s neck linker from 14 to 17 residues reduced both the run length and processivity (15). Thus, altering neck linker length differentially affects the processivity of Kif18A compared to Kinesin-1 and Kinesin-2. Although we don’t yet understand the molecular basis of these differences, one possible explanation may be due to unique loops in Kinesin-8 family motor domains (22, 38) that allow extended dwell times at microtubule ends and promote depolymerization of microtubules by Kip3p. Microtubule structure at a growing end is curved (39), unlike the rigid straight tracks along the lattice, and many proteins have a higher affinity for the growing tip (40, 41). Therefore, it is possible that Kif18A’s affinity for the microtubule lattice is reduced compared to Kinesin-1 or 2 motors, a trait that may be exacerbated by Kif18A sNL1’s faster velocity. Also of note, our single molecule studies were performed with C-terminal truncated forms of Kif18A to observe the influence of neck linker length without the additional tail microtubule-binding site. However, the full length protein used in the cellular studies does have this extra tether, which has been shown to increase the processivity of Kif18A (20), and Kif18A sNL variants are able to accumulate on the longer microtubules making up peripheral K-fibers, indicating that this decrease in run length does not fully explain the cellular localization phenotype.

While our data indicate that Kif18A’s neck linker length is important for navigating K-fiber regions containing HURP, there are certainly additional obstacles that Kif18A must contend with on K-fibers. The TACC3-ch-TOG-clathrin complex forms inter-microtubule bridges within K-fibers, influences the number of microtubules in a K-fiber, and is necessary for K-fiber stabilization (37, 42). Eg5 and Kif15 are kinesin motors that stabilize the spindle and occupy the same microtubule-binding site as Kif18A (2). The dynein/dynactin complex, which also shares an overlapping binding site with kinesins (43), transports checkpoint silencing proteins like Mad2 and BubR1 towards the spindle poles along K-fibers (44). Near the K-fiber plus ends, the kinetochore protein complex is its own gauntlet of dense obstacles (45). K-fibers are undoubtedly crowded tracks, but here we provide evidence that Kif18A is uniquely adapted to this environment to reach and accumulate on K-fiber ends, a trait that is required for the timely suppression of microtubule dynamics and chromosome alignment, ultimately ensuring equal distribution of genetic material to the next cell generation.

## Materials and Methods

### Plasmids and siRNAs

GFP-Kif18A sNL variants were constructed by Gateway Cloning technology (Thermo Fisher Scientific) and Quikchange site-directed mutagenesis (Agilent) into a pCMV-eEGP-N1 backbone containing the full length WT Kif18A sequence with N-terminal GFP tag and C-terminal His tag in frame. Specific deletions for each construct are shown in Figure 1. For protein purification, WT Kif18A and sNL1 were recombined from a pCR8/GW/TOPO entry vector into a pET-eGFP destination vector with N-terminal GFP tag and C-terminal FLAG tag with an LR reaction (Thermo Fisher Scientific). Scrambled control siRNA obtained from Thermo Fisher Scientific (Silencer Negative Control #2). Kif18A siRNA anti-sense sequence (4, 11, 22): 5’-GCUGGAUUUCAUAAAGUGG −3’ (Ambion). HURP siRNA anti-sense sequence: 5’-AAGGAGUCCAGGUGUAACUGG-3’ (Qiagen).

### Cell culture and Transfections

HeLa cells were cultured at 37°C with 5% CO_2_ in MEM-alpha medium (Life Technologies) containing 10% FBS plus 1% penicillin/streptomycin (Gibco). For siRNA transfections in a 24 well format, approximately 80,000 cells in 500 pμ MEM-alpha medium were treated with 35 pmol siRNA and 1.5 pμ RNAiMax (Life Technologies), preincubated for 20 minutes in 62.5 pμ OptiMeM (Life Technologies). Cells were treated with siRNAs for 48 hours before fixing or imaging. 24 hours prior to fixation or imaging, the media was changed and plasmid transfections were conducted similarly, but with 375 ng Kif18A or HURP plasmid DNA and 2 pμ LTX (Life Technologies).

### Cell fixation and immunofluorescence

For fixation, HeLa cells were treated with 20 μM MG132 (Selleck Chemicals) for 1-2 hours before fixation. Cells were fixed on coverslips in ‐20°C methanol (Fisher Scientific) plus 1% paraformaldehyde (Electron Microscopy Sciences) for 10 minutes on ice, dunked briefly and then washed 3X for 5 minutes each in TBS. Coverslips were blocked with 20% goat serum in antibody-diluting buffer (Abdil: TBS pH 7.4, 1% bovine serum albumin, 0.1% Triton-X, and 0.1% sodium azide) for 1 hr at room temperature before the primary antibodies diluted in Abdil were added for 1 hour at room temperature: mouse anti-α tubulin at 1 μg/mL (Sigma-Aldrich), mouse anti-γ-tubulin at 1 μg/mL (Sigma-Aldrich), rabbit anti-GFP at 4 μg/mL (Molecular Probes), rabbit anti-HURP at 2 μg/mL (Bethyl). Primary antibodies against human anti-centromere protein (ACA) (Antibodies Incorporated) were used at 2.5 μg/mL overnight at 4°C. Incubation with 1 μg/mL goat secondary antibodies against mouse, rabbit, or human IgG conjugated to Alex Fluor 488, 594, or 647 (Molecular Probes) occurred for 1 hour at room temperature. 2X TBS 5 minute washes were performed between rounds of primary and secondary antibody incubations, finishing with 3X TBS washes before mounting with Prolong Gold anti-fade mounting medium with 4’,6-Diamidine-2’-phenylindole dihydrochloride (DAPI) (Molecular Probes).

### Microscopy

Cells were imaged on a Nikon Ti-E inverted microscope (Nikon Instruments) with Nikon objectives Plan Apo 40X DIC M N2 0.95 NA, Plan Apo λ 60X 1.42NA, and APO 100X 1.49 NA with a Spectra-X light engine (Lumencore) and environment chamber. The following cameras were used: Clara cooled-CCD camera (Andor) and iXon X3 EMCCD camera (Andor). The Nikon Ti-E microscope is driven by NIS Elements software (Nikon Instruments). Live cell imaging is described in SI methods. TIRF microscopy was performed at room temperature using an inverted Eclipse Ti-E microscope (Nikon) with a 100x Apo TIRF objective lens (1.49 N.A.) and dual iXon Ultra Electron Multiplying CCD cameras, running NIS Elements version 4.51.01. Rhodamine-labeled microtubules were excited with a 561 and a 590/50 filter. WT KIF18A Δ480-GFP or SNL2 KIF18AΔ480-GFP motors were excited with a 488 laser and a 525/50 filter. Alexa 647 labeled tau was excited with a 640 laser and 655 filter. All movies were recorded with an acquisition time of 100 msec.

### Kif18A line scan analysis

Fixed and stained cells were imaged at 100X with 0.2 μm z-stacks taken through the full cell. Line scans were manually measured using the α tubulin and ACA fluorescent intensities to identify well-defined and locally isolated K-fibers. Lines we subdivided into central or peripheral spindle K-fiber classification based on their location in the spindle; ambiguous regions were avoided. The profile intensities for α tubulin, ACA, and GFP-Kif18A were measured and recorded. Each channel was normalized internally to its highest value. Line scans were aligned in block by peak ACA intensity, and averaged for each pixel distance. Standard deviations are reported. Statistical comparisons were performed by fitting a linear regression of the accumulation slope (−0.8 μm to 0 μm) and comparing each sNL variant to that of WT Kif18A at central and peripheral K-fibers using an F test. To determine the ratio of tip to lattice Kif18A fluorescent intensity, line scans in the central spindle were normalized and aligned to peak ACA intensity as above, but then each line was sectioned between ‐0.768 and ‐0.32 μm (lattice) and ‐0.32 to 0.0 μm (tip) and the area under the curve calculated for each. These area under the curve calculations were then divided to determine the ratio of tip to lattice Kif18A accumulation for each individual line scan, which was then graphed for each condition.

### Chromosome alignment analysis

Quantification of chromosome alignment was performed as described previously (11, 22). Briefly, single focal planes were imaged of metaphase cells with both spindle poles in focus. The Plot Profile command in ImageJ was used to measure the distribution of ACA-labeled kinetochore fluorescence within a region of interest (ROI) defined by the length of the spindle with a set height of 17.5 μm. The ACA signal intensity within the ROI was averaged along each pixel column, normalized, and plotted as a function of distance along the normalized spindle pole axis. These plots were analyzed by Gaussian fits using Igor Pro (Wavemetrics). The full width at half maximum intensity (FWHM) for the Gaussian fit and the spindle length are reported for each cell analyzed.

### Single molecule TIRF motility assays with and without tau

Kif18A protein purification is described in the SI methods. Bovine tubulin and tau (3RS) was purified and tau labeled with Alexa 647 as reported previously (46). Rigor kinesin motility assays are described in the SI methods. For motility assays with or without tau, flow chambers used in *in vitro* TIRF experiments were constructed as previously described (46). Briefly, flow chambers were incubated with monoclonal anti-β III (neuronal) antibodies (Sigma-Aldrich) at 33 μg/ml for 10 minutes, then washed twice with 0.5 mg/ml bovine serum albumin (BSA; Sigma Aldrich) and incubated for 2 minutes. 1 μM of microtubules (all experimental conditions) were administered and incubated for 10 minutes. Non-adherent microtubules were removed with a BRB80 wash (80 mM PIPES, 1mM MgCl_2_, 1 mM EGTA, 10mM DTT, 1mM MgCl_2_, and an oxygen scavenger system [5.8 mg/ml glucose, 0.045 mg/ml catalase, and 0.067 mg/ml glucose oxidase; Sigma Aldrich]) supplemented with 20 μM paclitaxel (Sigma-Aldrich) and 100 mM KCl. Kinesin motors in BRB80 + 100 mM KCl and 1 mM ATP were added to the flow cell just before image acquisition. For rhodamine-labeled microtubules, unlabeled GTP tubulin was mixed with rhodamine-labeled tubulin (Cytoskeleton) at a ratio of 100:1. The tubulin mixture was polymerized at 37°C for 20 minutes and stabilized with 20 μM paclitaxel in DMSO. For experiments containing tau, tubulin polymerization was performed as above but in the absence of rhodamine-labeled tubulin. Polymerized microtubules were incubated with 25 nM Alexa 647 labeled tau (1:40 tau to tubulin ratio) for an additional 20 minutes at 37°C. Tau and microtubules mixture was centrifuged at room temperature for 30 minutes at 15,000 rpm. The pellet was then resuspended in BRB80 + 100 mM KCl and 20 μM taxol solution. It should be noted that the tau and rigor kinesin single molecule assays were performed in different buffers (BRB80 + 100 mM KCl vs. MAB) resulting in slight variation of Kif18A velocities and run lengths between the data sets.

### TIRF assay motility analysis

Motility events were analyzed as previously reported (16, 18). In brief, overall run length motility data was measured using the ImageJ (NIH) MTrackJ plug-in, for a frame-by-frame quantification of Kif18A motility. Characteristic and corrected run lengths and track lengths were calculated as previously described (47). Average run length and velocity were plotted as a histogram with mean and SD. Statistical significance was determined by unpaired t-test. Kymographs of motor motility were created using the MultipleKymograph ImageJ plug-in, with a line thickness of 3.

## Acknowledgments

We wish to thank Puck Ohi for his thoughtful discussion and generous gift of the GFP-HURP plasmid created by Sophia Gayek, and Lynn Chrin for her work purifying and labeling 3RS tau and rigor kinesin. We also thank Maria Kogan for writing a custom Matlab program for line scan analysis. This work was funded by NIH NIGMS R01GM121491 awarded to J. Stumpff. H. Malaby is currently supported by a Department of Defense PRCRP Horizon Award (W81XWH-17-1-0371). The authors declare no competing financial interests.

## Abbreviations

(WT): Wild-type
(sNL): short neck linker
(K-fiber): Kinetochore Microtubule bundle
(ACA): Human centromere A

